# Multiscale imaging of corneal endothelium damage and effects of Rho Kinase inhibitor application in mouse models of acute ocular hypertension

**DOI:** 10.1101/2023.05.18.541299

**Authors:** Zhen Cai, Yang Zhang, Raymond S. Fang, Benjamin Brenner, Junghun Kweon, Cheng Sun, Jeffery Goldberg, Hao F. Zhang

**Author notes:** These authors contributed equally to this work (Zhen Cai, Yang Zhang).

## Abstract

We developed a multiscale optical imaging workflow, integrating and correlating visible-light optical coherence tomography, confocal laser scanning microscopy, and single-molecule localization microscopy to investigate the mouse cornea damages from the *in-vivo* tissue level to the nanoscopic single-molecule level. We used electron microscopy to validate the imaged nanoscopic structures. We imaged wild-type mice and mice with acute ocular hypertension and examined the effects of Rho Kinase inhibitor application. We defined four types of intercellular tight junction structures as healthy, compact, partially-distorted, and fully-distorted types by labeling the Zonula occludens-1 protein in the corneal endothelial cell layer. We correlated the statistics of the four types of tight junction structures with cornea thickness and intraocular pressure. We found that the population of fully-distorted tight junctions correlated well with the level of cornea edema, and applying Rho Kinase inhibitor reduced the population of fully-distorted tight junctions under acute ocular hypertension.

## Introduction

Biological systems coordinate multiple components within and across multiple length- and time-scales.^1-4^ In the eye, key anatomical components, including cornea^5^, iris, vitreous, lens, retina, and optic nerve head (**Fig. 1a**), collectively support its optical and neurological capabilities. The optically transparent cornea contains multiple layers, including the epithelial cell layer, endothelial cell layer, and acellular stromal layer (**Fig. 1b**.) The corneal epithelial and stromal layers constitute approximately one-third and two-thirds of the total thickness, and the corneal endothelial cells (CECs) only form a ∼2-μm thick monolayer in the cornea.^6^ Cornea dysfunction can lead to vision loss and blindness, affecting more than 12.7 million people globally.^4,7^ The corneal function is fundamentally powered by nanoscale biomolecular machinery, assemblies into subcellular organelles, and eventually forming specific cell types (e.g., endothelial and epithelial cells).^2,8-12^ Reversely, the morphological and physiological changes of the cornea as a whole can also affect its cellular and biomolecular activities. ^1,2,8-12^ Therefore, multiscale analyses, covering lengthscales from the *in-vivo* whole cornea anatomy to intracellular and intercellular single molecules, are important to comprehensively understand corneal health, diseases, and therapies.

**Fig. 1.**
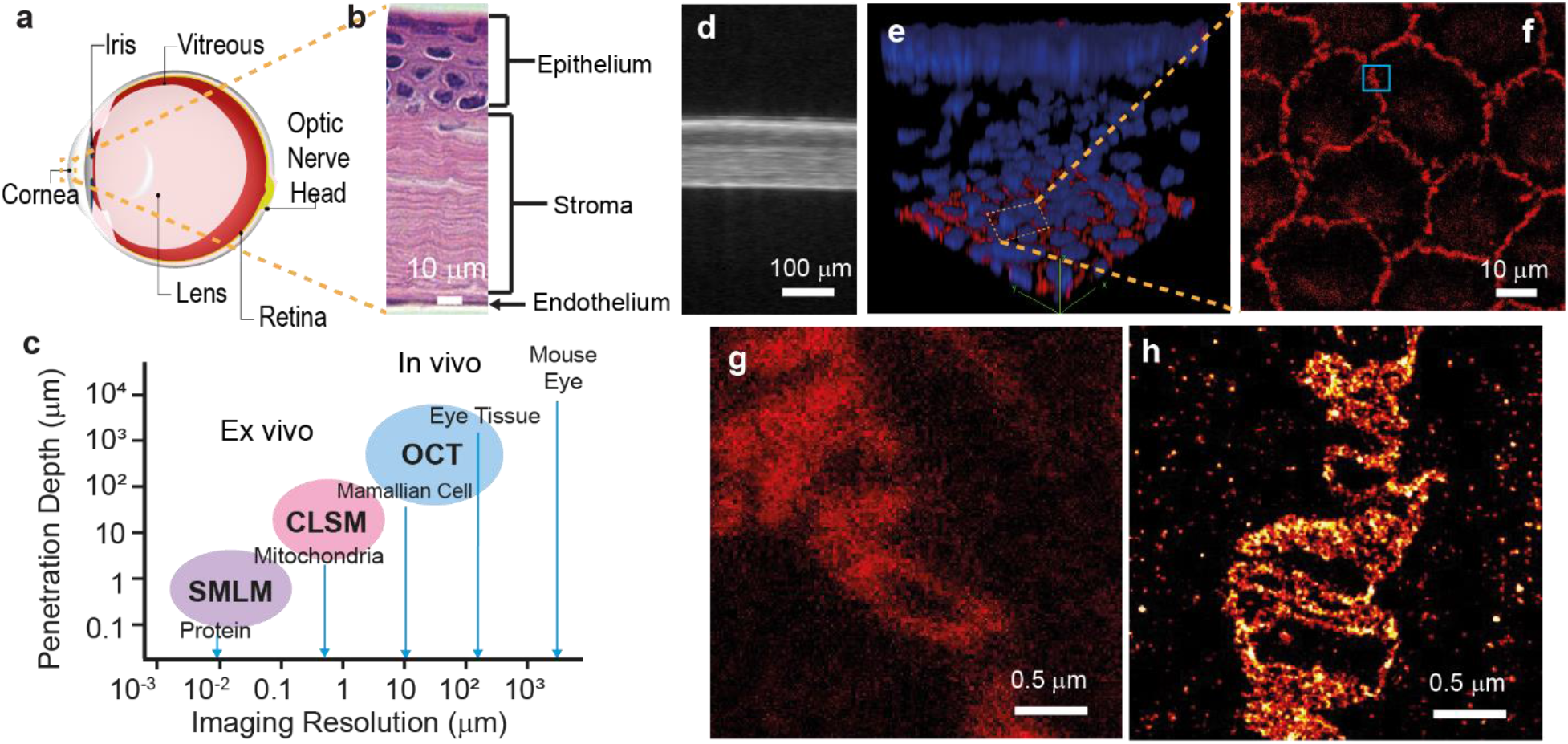
The concept of the multiscale imaging workflow. (a) Illustration of a mouse eye; (b) A histological image of mouse cornea; (c) Comparing imaging resolution and penetration depth among OCT, CLSM, and SMLM. (d) An *in-vivo* vis-OCT B-scan image of a wild-type mouse cornea; (e) A 3D visualization using CLSM of a fixed mouse cornea flat-mount immunofluorescent labeled with a tight junction marker (ZO-1, red) and nuclear stain (DAPI, blue); (f) An CLSM image of the CEC tight junctions showing the classical hexagonal boundaries; (g) A magnified view of the area highlighted in panel e; (h) An SMLM image, with a similar FOV as the panel g, visualizing the nanoscale structures and filaments of the ZO-1 organizations between adjacent cells.

To satisfy the need for multiscale analysis, we combined three optical imaging modalities with complementary resolutions and contrasts to achieve investigations from the *in-vivo* intact cornea to nanoscale intercellular structures (**Fig. 1c**). At the *in-vivo* intact cornea level, we applied noninvasive visible-light optical coherence tomography (vis-OCT), which can achieve an axial resolution of 1.3-μm and image the entire mouse anterior segment (**Fig. 1c** top right region),^13-15^ to monitor overall corneal health or damages in mice. At the *ex-vivo* cellular level, we applied confocal laser scanning microscopy (CLSM), which offers a lateral resolution of up to 200 nm and an imaging penetration depth of up to several hundred micrometers (**Fig. 1c** center region),^16^ to classify CECs in flat-mount corneal tissue samples. However, due to the optical diffraction limit, CLSM cannot resolve nanoscale intercellular structures.^17^ We recently developed a super-resolution optical imaging workflow to visualize, for the first time, the nanoscale subcellular structures in the corneal endothelium using single-molecule localization microscopy (SMLM).^18^ SMLM offers a spatial resolution of 20 nm at a penetration depth of a few micrometers (**Fig. 1c** bottom left region). ^19-22^ Hence, at the nanoscopic level, we applied SMLM to image intercellular nanostructures in CECs.

Acute ocular hypertension (AOH) and the progression of acute primary angle closure glaucoma (APACG) can cause wounds and dysfunctions in CECs, leading to vision loss.^23^ Recent studies found that adding Rho Kinase (ROCK) inhibitor to eye drops promoted corneal endothelial wound healing in different animal models^24-30^. ROCK signaling pathways contribute to various cellular functionalities, such as cell adhesion, motility, proliferation, differentiation, and apoptosis ^31,32^. The role of ROCK has been intensively investigated as a therapeutic target in a broad spectrum of diseases such as vascular disease, cancer, degenerative neuronal disease, asthma, and glaucoma ^24,26,27,29,30,33-36^. Using multiscale imaging, we also investigated the effect of ROCK-inhibitor on the overall corneal damage and intracellular damage among CECs in the mouse model of AOH.^37^ The mouse models of AOH have been broadly used to study various glaucoma damages in both the anterior and posterior segments, which induces significant changes in endothelial density, morphology, and nanoscale tight junction structures. At the subcellular level, the damages in CEC tight junctions could lead to an uncontrolled fluid passage from the anterior chamber, causing edema and stroma damage. ^38^

In this work, we established the workflow of multiscale cornea imaging in wild-type mice and quantified corneal and CEC damages and ROCK-inhibitor effects in mice with AOH. More specifically, we used vis-OCT to quantify the corneal anatomy *in vivo* (**Fig. 1d**); then, we used CLSM to image CEC’s morphological labeled by a tight junction Zonula occludens-1 (ZO-1) protein (**Figs. 1e-1f**) in flat-mount cornea samples; and finally, we used SMLM to quantify nanoscale morphological variations of ZO-1 labeled tight junctions and validated by scanning electron microscopy (SEM) in flat-mount cornea samples (**Fig. 1g**), where the tight junction nanostructures are not resolvable by CLSM (Fig. 1h).

## Methods and materials

**Fig. 2** shows our multiscale imaging workflow, from generating the mouse models to SMLM imaging. First, we treated the left eyes of young mice at 8-12 weeks of age with ROCK-inhibitor (ROCK group) or DMSO (control group) at four drops a day for 14 days. Then, we acquired *in vivo* vis-OCT images of left corneas before AOH in both groups. To introduce AOH, we cannulated the anterior chamber of the left eye to induce elevated IOP (∼40 mmHg) in the AOH groups for one hour. We also maintained 8 mmHg for one hour in the normal IOP groups. After a second round of *in vivo* imaging, we harvested the left corneas, fixed them, and immunofluorescent-labeled them to visualize ZO-1 or other intracellular structures of interest. Finally, we acquired CLSM, SMLM, and SEM images of the cornea samples for further analysis.

**Fig. 2.**
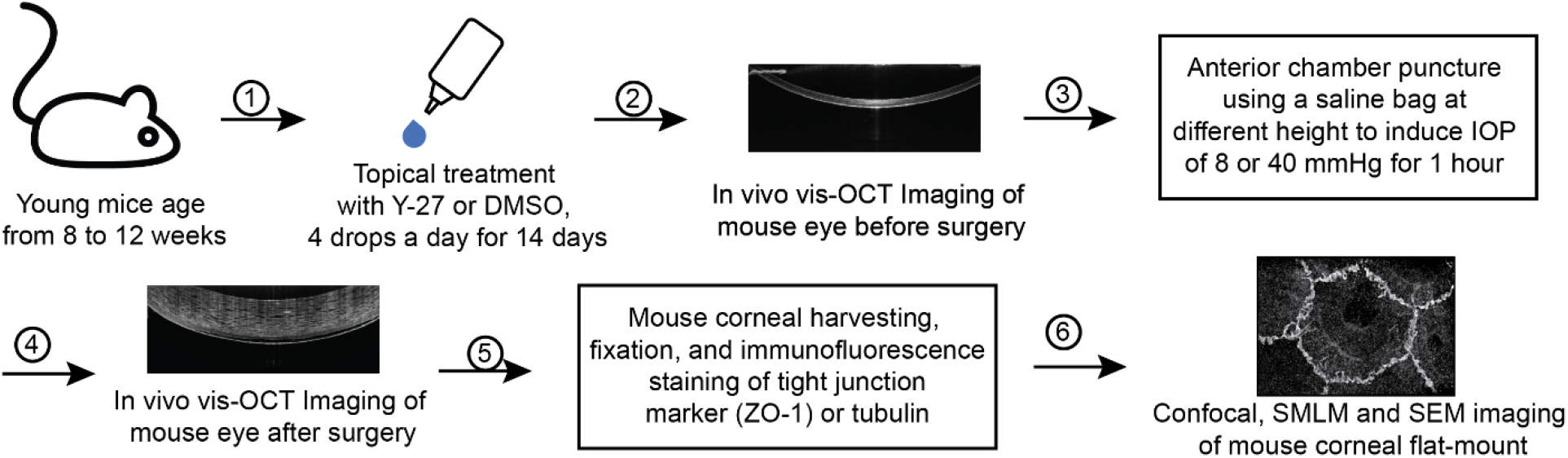
Experimental protocol for generating mouse models of AOH and subsequent application of ROCK inhibitor.

### AOH mice model generation

We used 8-12 weeks-old Wild-type C57BL/6 mice in our studies. The experimental animals were kept in the Center for Comparative Medicine at Northwestern University under normal lighting conditions with 12-h on and 12-h off cycles. All animal protocols were approved by the Institutional Animal Care and Use Committees of Northwestern University and conformed to the Association for Research in Vision and Ophthalmology Statement on Animal Research.

To create AOH, the mice were anesthetized via intraperitoneal injection (10 mL/kg body weight) of a ketamine/xylazine cocktail (ketamine: 11.45 mg/mL; xylazine: 1.7 mg/mL, in saline). We maintained the mouse body temperature within 34-36 °C using a rectal temperature probe (FHC Inc., catalog # 40-90-8D) with temperature feedback and its attached thermal heating pad. We used lidocaine as the lubricant for the temperature probe. We applied 1% Povidone-iodine for 60 seconds to the left eye and then washed the eyes with sterile PBS for conjunctival sac disinfection for 60 seconds. We inserted a 33-Gauge needle from the marginal cornea into the anterior chamber of the left eye. We checked the integrity of the iris under a surgical microscope before proceeding. Using manometry, we stabilized the needle with external mechanical support and controlled the IOP.^39^ More specifically, the eye was cannulated and connected to a syringe filled with PBS containing 5% mmol/L glucose. We controlled the IOP by adjusting the height of the syringe relative to the mouse eye. We elevated the syringe position to ∼55 cm above the mouse eye, corresponding to an IOP of around 40 mmHg, and maintained it for 1 hour. Afterward, we adjusted the syringe position to be ∼10 cm above the eye, corresponding to an IOP of 8 mmHg, and performed in vivo imaging. We documented the IOP every 15 minutes using a tonometer (iCare TONOLAB) and applied artificial tears (Alcon, NDC 17478-062-35) every 2 minutes. In addition, we checked the toe pinch reflex and respiratory rate every 15 minutes to ensure deep anesthesia.

### ROCK inhibitor application

To prepare the ROCK inhibitor solution, we first dissolved solid Y-27623 (Cayman Chemical 1000558310) in DMSO to achieve a concentration of 0.1 M and then diluted it with PBS to achieve a final concentration of 200 μM. For topical administration, we first immersed a sterile cotton tip applicator in the Y-27623 solution; then, we gently wiped the tip of the cotton tip applicator on the surface of the left cornea and repeated it four times with an interval of 10 minutes. In the control group, we applied a solution of 200 μM DMSO in PBS using the same method. We repeated all topical administrations daily for 14 days in both groups.

### *In vivo* vis-OCT imaging

We imaged the cornea using a custom-built vis-OCT system, operating between 510 nm and 610 nm as previously reported ^39-42^. The vis-OCT used an incident power of 1.5 mW on the cornea, and our imaging field of view was 2.7 × 2.7 mm^2^, containing 1024 × 1024 A-lines. We conducted *in vivo* vis-OCT imaging immediately after the AOH procedure without removing the needle. We applied artificial tears every 2 minutes during data acquisition to prevent corneal dehydration. Before measuring corneal thickness, we manually segmented each anatomical layer in the central corneal region. Four members independently measured the thickness of each corneal anatomical layer using ImageJ line profile measurement, and we reported the mean values in this work.

### Corneal sample preparation for CLSM and SMLM

After vis-OCT imaging, we removed the needle, euthanized the mice by cervical dislocation, and retrieved the cornea immediately. We first rinsed the cornea with PBS to remove any blood, pigment, and tissue residues attached to the cornea. Then, we cut the cornea into four equal pieces and fixed them in 0.5% Paraformaldehyde (PFA) at 25°C for 30 mins. The cornea samples were permeabilized with 1% Triton X-100 in PBS at 25°C for 30 min and blocked in 2% normal goat serum, 2% bovine serum albumin (BSA), and 1% Triton 100-X in PBS at 25°C for 2 hours. The cornea samples were further incubated with a primary antibody targeting a tight junction marker protein (ZO-1, #40-2200, Invitrogen) at 4°C for 48 hours. Lastly, the cornea samples were incubated with a secondary antibody (anti-rabbit Alexa Fluor 647, Thermofisher) at 4°C for 12 hours. We rinsed the cornea samples three times for 20 min at 25°C after each fixation, primary antibody incubation, and secondary antibody incubation step.

### SMLM imaging

After immunostaining, we placed the cornea samples on a rectangular No. 1.5 cover glass (22 × 60 mm^2^) with the CECs facing down according to the physiological curvature of the cornea and applied a few strips of double-sided tapes on the cover glass around the cornea samples. Next, we gently flattened the cornea sample using ophthalmic micro forceps, removed excessive PBS around the tissue samples, and covered the cornea samples with a No. 1 square cover glass (22 mm) to create an imaging chamber together with the double-sided tapes. Finally, we fill the imaging chamber with an appropriate imaging buffer solution for SMLM.

Before each imaging, we freshly prepared the imaging buffer solution containing 50 mM Tris (pH =8.0), 10 mM NaCl, 0.5 mg/mL glucose oxidase (Sigma, G2133), 2000 U/mL catalase (Sigma, C30), 10% (w/v) D-glucose, and 100 mM cysteamine. We added ∼10 μL of the imaging buffer to fill the imaging chamber and sealed it with black nail polish to prevent leaking during the imaging process. We refilled the imaging buffer every two hours in prolonged SMLM data acquisition sessions. Finally, we mounted the imaging chamber onto the SMLM microscope with the rectangular cover glass facing downward.

For SMLM imaging acquisition, we used a Nikon Ti-2E fluorescence microscope with a 100× objective lens (Nikon SR HP Apo TIRF, 100×) and a 647-nm continuous-wave laser to illuminate the sample using a highly inclined and laminated optical sheet (HILO) illumination with an angle slightly smaller than the critical angle at a power density of about 5 kW cm^-2^, as previously reported.^18,43-45^. The exposure time of each SMLM frame was 10 ms, and we acquired 30,000 frames from each sample for image reconstruction using ThunderSTORM plug-in of ImageJ.^46^

### CLSM imaging

We mounted the cornea samples in the same manner as we did for SMLM imaging. However, instead of the aforementioned imaging buffer solution, we used a PBS solution containing 1-μM Hoechst 33328 to fill the imaging chamber and counterstain the cell nucleus. We used a Leica SP5 confocal microscope to acquire all our CLSM images using a 20× oil immersion objective lens with a pixel size of 1 μm except **Fig. 1g** using a 60× oil immersion with a pixel size of ∼300 nm.

### SEM imaging

We first fixed the cornea samples in a solution containing 2.5% glutaraldehyde and 4% PFA in a 0.1M sodium cacodylate buffer pH 7.4. Then we fixed the corneal samples further with 1% osmium tetroxide and dehydrated them in a graded series of ethanol (30%, 50%, 70%, 85%, 95%) before critical point drying in a Tousimis Samdri-795 critical point drier. Before SEM imaging, we mounted the processed corneas to an aluminum SEM stub with carbon tape and silver paint and sputter-coated with 10 nm of Au/Pd in a Denton Desk IV sputter coater. Finally, we imaged all samples using a Hitachi SU8030 cFEG SEM at 10 kV.

## Results

### *In vivo* vis-OCT imaging measured bulk anatomical alterations in mouse corneas

**Fig. 3** and **Table 1** show the *in vivo* vis-OCT^39^ imaging of the corneas to assess the effects of AOH and ROCK inhibitor application. We compared the central corneal thickness of four groups: group C1 with normal IOP and DMSO treatment, group C2 with AOH and DMSO treatment, group C3 with normal IOP and Y-27623 treatment, and group C4 with AOH and Y-27623 treatment. **Figs. 3a-3d**, respectively show representative vis-OCT images from each of the four groups. In the control group C1 (**Fig. 3a**), vis-OCT well resolved the epithelium, stroma, and endothelium layers thanks to its 1.3-μm axial resolution,^47^ making it possible to measure variations in the thickness of these layers. **Fig. 3b** shows that AOH significantly increased stromal thickness with characteristic edema in group C2 compared with group C1. **Fig. 3c** shows that topical administration of Y-27623 in group C3 induced no change in corneal thickness compared to the control group C1. The cornea treated with Y-27623 (**Fig. 3d**) in group C4 exhibited less severe edema and a minor increase in stromal thickness compared with the untreated cornea subjected to AOH in group C2 (**Fig. 3b**).

**Table 1.** *In vivo* thickness measurements by vis-OCT (± denotes standard deviations.)

|  | Control | AOH | ROCK | ROCK+AOH |
| --- | --- | --- | --- | --- |
| Cornea thickness ($\mu\text{m}$)<sup>a</sup> | 104.2±11.7 | 444.6±71.6 | 89.3±17.7 | 321.9±66.9 |
| Endothelium ($\mu\text{m}$) | 4.5±1.9 | 4.3±1.6 | 6.9±2.9 | 7.7±1.6 |
| Stroma ($\mu\text{m}$) | 64.2±8.9 | 376.9±57.4 | 55.5±15.8 | 285.5±66.8 |
| Epithelium($\mu\text{m}$) | 37.4±6.4 | 63.9±18.0 | 31.5±3.3 | 38.9±3.4 |

**Fig. 3.**
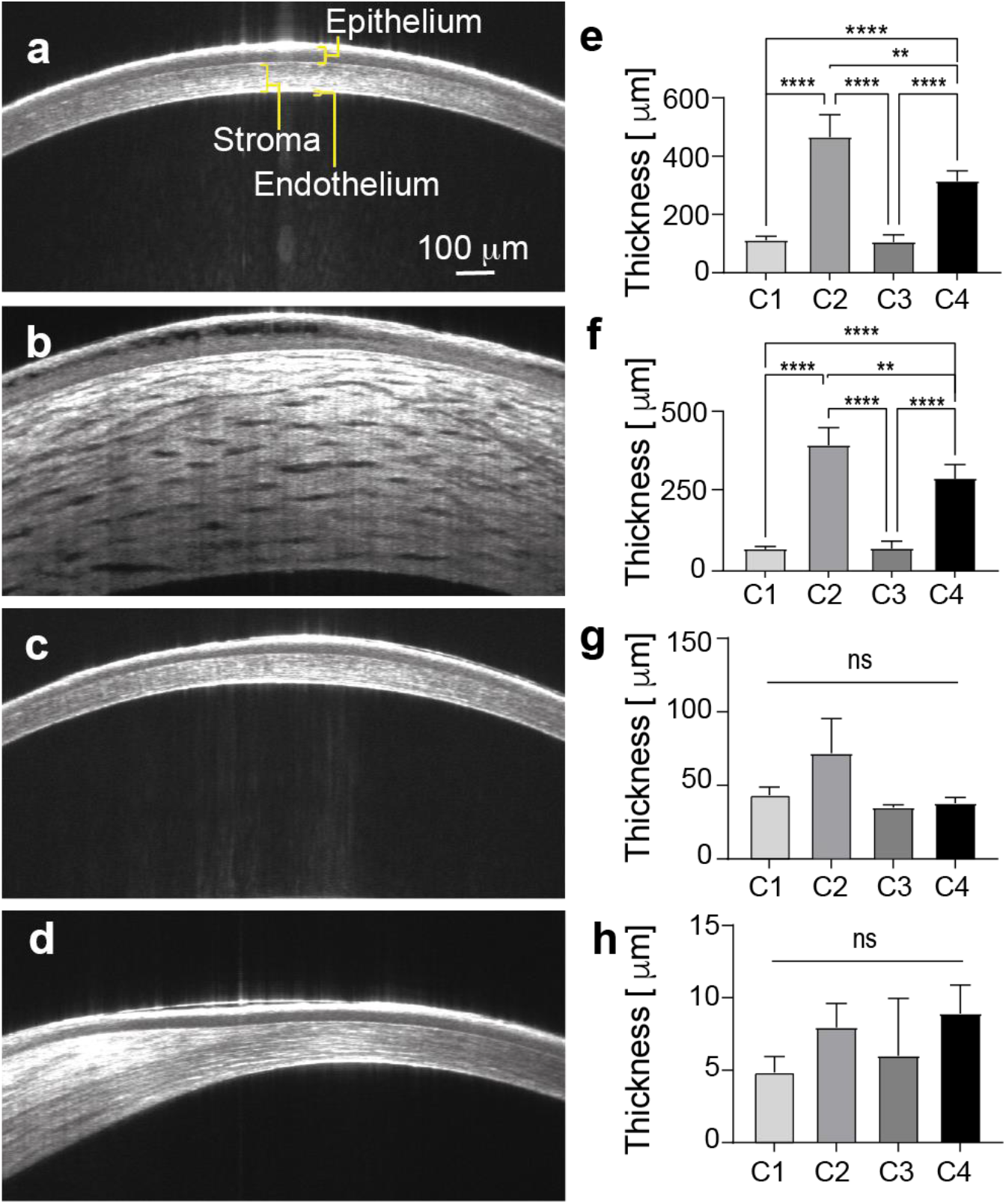
*In vivo* vis-OCT image of mouse cornea at four different conditions. (a) Condition C1 with normal IOP and DMSO treatment; (b) Condition C2 with elevated IOP and DMSO treatment; (c) Condition C3 with normal IOP and Y-27623 treatment; (d) Condition C4 with elevated IOP and Y-27623 treatment; (e) total corneal thickness variation; (f) stromal thickness variation; (g) corneal epithelium thickness variation; (h) corneal endothelium thickness variation. N=3 for each condition.

Next, we compared vis-OCT-measured thickness variations of the total cornea, epithelium, stroma, and endothelium in the four groups, as shown in **Table 1** and **Figs. 3e-3h**. In the control group C1, the corneal epithelium, stroma, and endothelium have average thicknesses of 37.4 μm, 64.2 μm, and 4.5 μm, respectively, and the average total corneal thickness is 104.2 μm. Compared with the C1 group, the total corneal thickness increased 4-fold and 3-fold in group C2 and group C4, respectively, (**Fig. 3e**), and stromal thicknesses increased 5-fold and 4-fold in the C2 and C4 groups, respectively (**Fig. 3f**). The reductions in the total cornea and stroma thicknesses are statistically significant between the C2 and C4 groups, suggesting topical administration of Y-27623 reduced edema in corneas subjected to AOH. However, we did not observe statistically significant changes in the endothelium (**Fig. 3g**) and epithelium (**Fig. 3h**) thicknesses in groups C2 and C4 compared with group C1. In addition, we did not observe statistically significant thickness changes in the total cornea, epithelium, stroma, and endothelium between the groups C1 and C3, suggesting no effect from the topical administration of DMSO.

### CLSM imaging classified CECs based on different ZO-1 morphologies

After *in-vivo* vis-OCT imaging, we harvested the corneas and examined the CEC morphological changes in the four groups. Multiple studies reported the influence of external conditions (*e*.*g*., elevated IOP^26,28^, perfusion with diamide^48^) on cell-cell junctions in CECs that play key roles in endothelial permeability. **Fig. 4** shows representative CLSM images of CECs in groups C1-C4, where the cell nuclei are pseudo-colored in blue, and the labeled ZO-1 tight junctions are pseudo-colored in red. Tile-scan epifluorescence microscopy images are shown in Supplementary **Figs. S1-S4**. Although previous studies observed tight junction disruption in rat corneas under AOH^37^, we categorized the tight junction morphologies into four different types from CLSM images, which we further correlated with vis-OCT and SMLM findings.

**Fig. 4.**
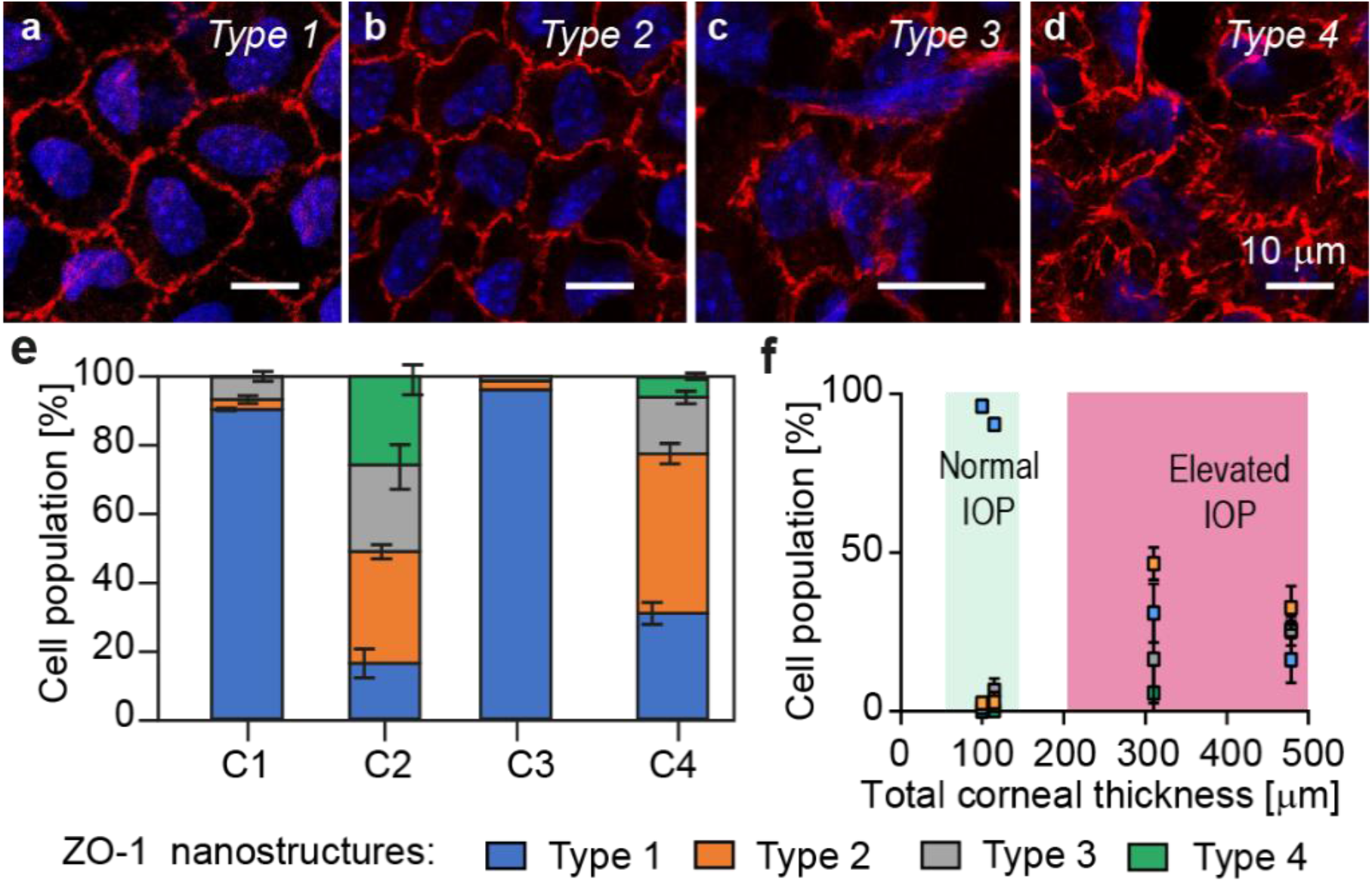
Representative CLSM images of a cluster of CECs (a) normal Type 1 ZO-1 structures; (b) compact Type 2 ZO-1 structures; (c) partially-distorted Type 3 ZO-1 structures; (d) fully-distorted Type 4 ZO-1 structures; (e) statistics of cell population exhibiting the four types of ZO-1 structures among 500-1500 cells from three mice; (f) scatterplot of cell population measured from CLSM versus total corneal thickness measured from vis-OCT; Error bar: standard deviation; n=4.

In a healthy cornea, ZO-1 is known to form regular hexagonal or pentagonal patterns in CECs and is organized continuously in adjacent cell borders, which we define as Type 1 structure (**Fig. 4a**). **Fig. 4b** shows that ZO-1 can still form the hexagonal or pentagonal patterns but exhibits more compact and slightly torturous boundaries among adjacent cells, which we define as Type 2 structure. **Fig. 4c** shows highly torturous and discontinuous ZO-1 structures with partial distortion among adjacent cells, which we define as Type 3 structure. Lastly, **Fig. 4d** shows ZO-1 structures severely pulled apart among the adjacent cell border with ZO-1 nanofibers stretched from the cell borders, which we define as Type 4 structure.

Based on the four defined ZO-1 morphological types, we obtained their respective statistical distributions in groups C1-C4. We imaged 500-2000 CECs from three mice in each group using epifluorescence microscopy with tile-scan function (Ti2E, Nikon). Four lab members independently classified ZO-1 structures based on the definition described above. **Fig. 4e** and **Table 2** show the percentage populations of the four CECs morphology types. Among all the CECs ZO-1 labeled cell junctions from the groups C1 and C3, 90%-95% are Type 1, 3% are Type 2, and <0.2% are Type 4. We found 6.6% and 1.2% Type 3 CECs in groups C1 and C3, respectively. In group C2, the Type 1 CECs population dropped to 16%, with an increase of Type 2, Type 3, and Type 4 CECs to 32.6%, 25.3%, and 25.8%, respectively. In group C4, 30.92% CECs remained healthy (Type 1), and the populations for Type 2, Type 3, and Type 4 CECs changed to 40.51%, 16.49%, and 5.83%, respectively. Compared with group C2, higher populations of Type 1 and Typ2 CECs in the Y-27623 treated group C4 contributed to the less severe edema in the stroma, agreeing with the *in vivo* thickness measurements by vis-OCT. **Fig. 4f** shows the percentage populations of the four types of CECs with respect to the corresponding total corneal thickness. When the total corneal thickness increased, the percentage population of the normal Type 1 CECs (blue squares) dropped, and the percentage populations of the Types 2-4 CECs increased.

**Table 2.** Percentage population of the four types of ZO-1 structures in CECs imaged by CLSM.

|  | Type 1 | Type 2 | Type 3 | Type 4 |
| --- | --- | --- | --- | --- |
| **C1** | 91.10±3.15% | 3.59±1.71% | 5.25±3.77% | 0.14±0.20% |
| **C2** | 15.81±5.10% | 32.61±9.32% | 28.33±13.90% | 25.81±9.61% |
| **C3** | 96.56±0.97% | 2.63±0.45% | 0.80±0.93% | 0.02±0.03% |
| **C4** | 29.86±6.84% | 50.28±7.31% | 28.33±4.59% | 4.18±2.81% |

### SMLM visualization of the four types of the ZO-1 nanostructures

Although we qualitatively identified four types of CECs with distinct ZO-1 structures, the exact ZO-1 morphology at the nanoscopic scale remains unresolvable by CLSM due to its diffraction-limited spatial resolution (**Fig. 1g**). To quantitatively define the ZO-1 nanostructure in these four types, we applied SMLM to both visualize and measure the with of ZO-1 labeled intercellular CEC junctions. Compared with CLSM imaging, we focused on a much smaller FOV in SMLM.

In Type 1 ZO-1 structures (**Fig. 5a**), we observed unique nanoscopic organizations of tight junctions organized along the boundaries between adjacent CECs with an average width of 1.3 μm (highlighted by solid yellow lines). We also observed the discontinuity of the ZO-1 pattern at the Y-junction where the three cells intersect, a known nanoscale feature of tight junctions associated with the relatively low-stress resistance of corneal endothelium^49^ (highlighted by the green arrow in **Fig. 5a**). Interestingly, we also observed fine discontinued nanoscale pores along the cell-cell borders, as the blue arrow indicated area with a pore size of ∼600 nm^2^. For the Type 2 ZO-1 structures, SMLM (**Fig. 5b**) shows compact tight junctions with an average width of 0.4 μm. The discontinuity of these ZO-1 structures is reduced or disappeared at the Y-junctions and along the cell borders (green arrows in **Fig. 5b**) compared to the Type 1 ZO-1 structure. In a typical Type 3 ZO-1 structure, elevated discontinuity boundaries among adjacent cells with distorted fiber branches are visualized (**Fig. 5c**), where the non-distorted portion of the ZO-1 structure width is comparable to the Type 2 ZO-1 structure. Lastly, the Type 4 ZO-1 structure shows complete separations of the tight junctions between adjacent cells along the cell borders and at the Y-junctions, with individual junction fibers clearly visualized (**Fig. 5d**).

**Figure 5.**
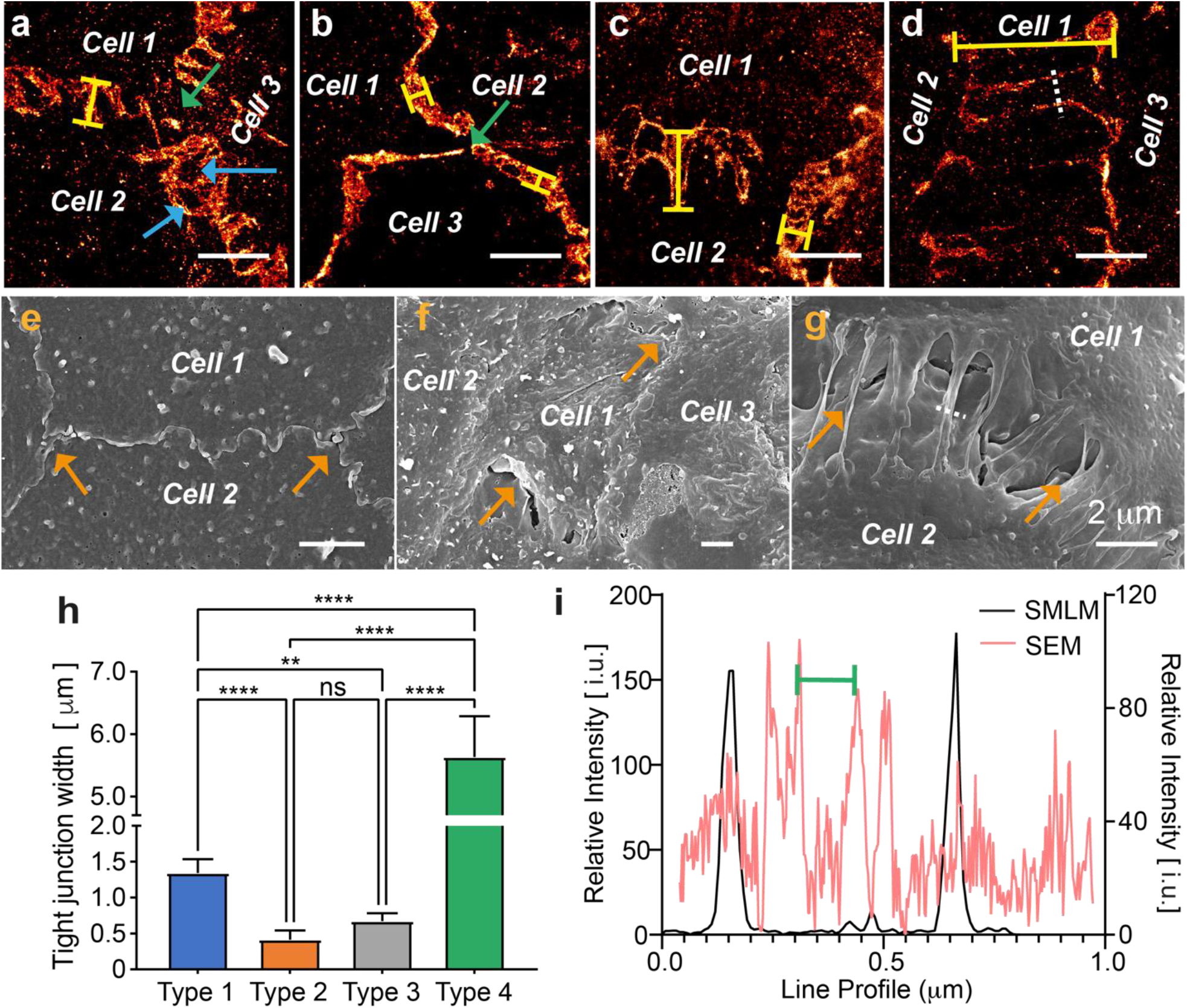
(a-d) SMLM images of (a) Type 1, (b) Type 2, (c) Type 3, and (d) Type 4 ZO-1 structures in CECs. The solid yellow lines in each panel illustrate the unambiguous thickness measurement of tight junction thickness using SMLM; The green and yellow arrow-pointed areas indicate the classical tight junction discoutinuities at three-cell Y-junctions and newly-discovered tight junction pores using SMLM. (e-g) SEM image of CECs with (e) healthy tight junctions; (f) partially damaged tight junctions; (g) fully-distored tight junctions. (h) Tight junction widths measured from SMLM from the four types of CECs from the solid yellow lines. N = 5; (i) Line profiles of the stretched intercellular space ZO-1 nanofiber from the positions highlighted by the white dashed lines in panels d and g.

To further validate SMLM-imaged ZO-1 nanostructures, we imaged the four types of ZO-1 structures using SEM. Due to SEM’s imaging contrast limit, we could only visualize the superficial morphologies of the CECs without any specificity to ZO-1 proteins. From the SEM images, we examined how tightly adjacent cells are bonded in the four types of CECs. **Fig. 5e** shows similar cell boundaries that match the normal (Type 1) or compact (Type 2) ZO-1 structures shown in **Figs. 5a-5b**. SEM images of Type 3 CECs show similar “wrinkled” boundaries and partial separations between cells, as shown in **Fig. 5f**. For the Type 4 ZO-1 structure, SEM captured the stretched junction fibers when tight junctions were separated by the elevated IOP (**Fig. 5g**), closely matching the SMLM image shown in **Fig. 5d**.

We compared the SMLM measured ZO-1 structure widths among the four CEC types in **Fig. 5h** and **Table 3**, where the averaged widths of the four types are 1.3 μm, 0.4 μm, 0.6 μm, and 8.9 μm, respectively. Except for the ZO-1 structures between Type 2 and Type 3 CECs, all other ZO-1 structure types showed statistically significant differences in width. We also compared the sizes of individual junction fibers in Type 4 CECs imaged by SMLM and SEM (**Fig. 5i**) as the intensity profiles from the positions highlighted by the white dashed lines in **Figs. 5d & 5g**. Particularly, the full-width-at-half-maximum (FWHM) fiber size is ∼ 50 nm in SMLM, while those from the SEM measurements (inner-membrane distance as indicated by the green scale in **Fig. 5i**) were ∼70 nm. In summary, these nanoscale features visualized by both SMLM and SEM consistently showed a significantly increased ZO-1 width among CECs subjected to elevated IOP.

**Table 3.** SMLM measured widths of the four types of ZO-1 structures in CECs.

|  | Type 1 | Type 2 | Type 3 | Type 4 |
| --- | --- | --- | --- | --- |
| **Width ($\mu\text{m}$)** | 1.3±0.2 | 0.4±0.1 | 0.7±0.1 | 5.6±0.7 |

## Discussion

In this work, we report a multiscale workflow to image mouse cornea from *in-vivo* tissue level, including corneal thickness, to *ex-vivo* single-molecule level, including nanoscale morphological changes in the ZO-1 labeled CEC tight junctions. Correlating OCT and fluorescence microscopy has been widely used to study cellular and subcellular damage associated with eye diseases.^13^ However, the spatial resolution of conventional fluorescence microscopy technologies is fundamentally constrained by the optical diffraction limit, which prevents the visualization of underlying nanoscopic and single-molecule processes. SMLM overcomes the diffraction limit and offers a resolution down to ∼10 nm.^20^ SMLM requires stringent sample preparation protocols to preserve the nanoscale ultrastructure and minimize non-specific single-molecule labeling, which might not be detected in conventional fluorescence microscopy because of SMLM’s single-molecule detection sensitivity. We investigated the concentration, temperature, and incubation time in the fixation and antibody labeling steps, respectively, as detailed in the Method Section, to obtain the optimal sample preparation conditions.^18^

This new multiscale imaging workflow also allowed the first correlation studies of clinically-relevant damages in the mouse cornea. First, we revealed the nanoscopic variations of ZO-1 labeled tight junctions under the influence of AOH and subsequent ROCK inhibitor protection, as shown in **Fig. 5**. Under AOH, the observed tight junction nanofibers among adjacent cells in **Fig. 5d&5g** are likely attributed to the tunneling membrane nanotubes (TMT) based on their closely resembled morphologies compared with literature reports. More specifically, TMTs are defined as membrane protrusions with thicknesses of a few hundred nanometers and lengths at the micrometer scale, which link distant cell bodies. ^50,51,26^ TMT-like structures have been found in diverse mammalian cells, such as retinal pigment epithelial cells, dendritic cells in the corneal stroma, neuroblastoma cells, and breast cancer cells.^50^ However, to the best of our knowledge, this is the first report to visualize the formation of TMT in native tissue at the nanoscopic level by both SMLM and SEM. TMTs in the corneal endothelium under AOH, presumably function to alter Ca^2+^ signal transduction and other biomolecules in extended ranges of signal communication so that the cell-cell communication is maintained without using classical gap junction-mediated short-range signaling. We believe the nanoscale Type 4 ZO-1 structures (< 100 nm fibers) are not similar to microfilaments observed after corneal perfusion with diamide.^48^ Those microfilaments are formed on the edges among different clusters of CECs induced after corneal perfuion^45^ while the TMT is observed between individual CECs (**Fig. 5g**). However, we also note that the typical TMT forms between the gap junctions other than tight junctions where ZO-1 primarily highlights the latter.^50^ One hypothesis is that ZO-1 is mobile and can bind to stably arranged gap junction plaques, thus contributing to the formation of TMT.^52^

Our newly developed multiscale imaging workflow allowed us to demonstrate that applying a ROCK inhibitor can reduce the population of severely damaged ZO-1 structures, reducing the severity of corneal edema in mouse models subjected to AOH (**Figs. 3-5**). Additionally, our results support the hypothesis that ROCK inhibitor Y-27632 therapy may protect CECs from AOH, consistent with a previous report that found Y-27632 treatment helped maintain corneal clarity following APACG therapy^53^ by increasing aqueous outflow and reducing pressure on the optic nerve head.^54^ More specifically, our multiscale imaging workflow provides new insights into the subcellular alterations induced by ROCK inhibitors in maintaining corneal clarity, including protecting the morphologies of tight junctions in CECs. However, the detailed molecular pathways involved in AOH conditions and ROCK inhibitor therapy remain open questions.

There are several limitations in SMLM imaging for mouse ocular tissue. SMLM typically has a limited imaging depth of the field of ∼ 1-2 μm and an imaging penetration depth of ∼10 μm.^55^ This is because SMLM requires the collection of precious photons (<10^4^) emitted from the individual fluorescent labels with minimal optical attenuations and aberrations to localize single-molecule locations at nanometer precision. When applying SMLM to image tissue samples, only the biological structures that are close (< 10 μm) to the interface between the cover glass and the tissue sample can be routinely imaged. Imaging beyond a few μm range into the cornea is challenging due to the elevated autofluorescence background and the reabsorption and scattering of the emitted single-molecule fluorescence signal. Further technological developments to provide optical sections, brighter fluorescent labels, and imaging processing methods to analyze single-molecule emitters in deep ocular tissue could facilitate the visualization of other eye tissues at the nanoscale.

## Conclusion

We developed a multiscale corneal imaging workflow focusing on the corneal endothelium. For the first time, we revealed the alterations in tight junctions of the cornea endothelium at the nanoscopic level. Sub-100-nm ZO-1 tight junction fibers, likely the tunneling membrane nanotubes, perpendicular to the cell border, were observed among CECs. Vis-OCT-CLSM-SMLM correlation further suggests a close correlation between cornea and endothelium thicknesses and the percentile populations of four different CEC types. In the AOH model, the probabilities of CECs with the tight junction structure becoming more compact or stretched are 10-fold and >1000-fold higher than the healthy cornea. ROCK inhibitor application reduced the severely damaged CECs population by ∼80%. Detailed mechanisms inducing the stretched CECs and ROCK inhibitor effect in CECs under different levels of IOPs longitudinally should be further investigated.

## Acknowledgment

This work was supported in part by NIH grants R01EY029121, R01EY028304, U01EY033001, R01EY034740, R01GM139151, R01GM140478, T32GM142604, T32GM105538, R21GM141675, R01GM143397, and U54CA268084; NSF grants CHM-1954430 and EFMA-1830969; and a seed grant from Illinois Society for the Prevention of Blindness. We thank Northwestern Biological Imaging Facility (BIF), Mouse Histology and Phenotyping Lab (MHPL), Center of Advanced Molecular Imaging (CAMI), and Atomic and Nanoscale Characterization Experimental Center (NUANCE) core facilities for sample preparation and image acquisition support.

## Author contributions

Z.C., J.K. & R.F. generated mouse models, performed mouse surgery, and conducted vis-OCT imaging & analyses; Z.C., Y.Z. & B.B. performed SMLM imaging & analyses; Y.Z. performed CLSM and tile-scan fluorescence imaging; T.Z. & I.R performed additional imaging analyses. Z.C., Y.Z. & H.F.Z. conceived the study. J.G. & H.F.Z. supervised the project. All coauthors wrote the manuscript.

## Additional information

Supplementary information is available for this paper at http://doi.org/xxxxx.

