## Supplemental figures for "Multiscale imaging of corneal endothelium damage and effects of Rho Kinase inhibitor application in mouse models of acute ocular hypertension"

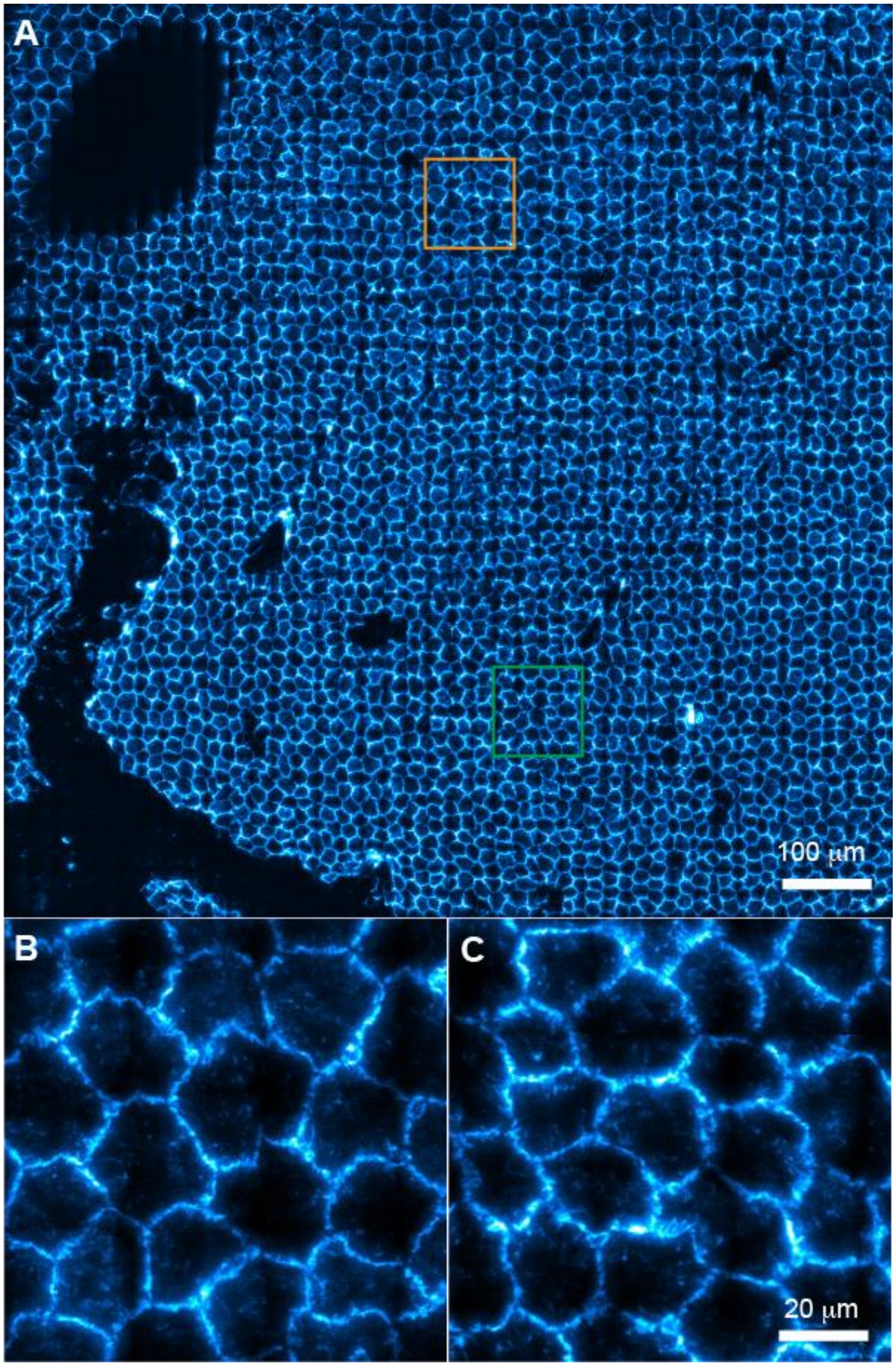


**Fig. S1.** (**A**) A representative tile-scan epifluorescence image with a scanning area of 1x1 $mm^2$ for the corneal endothelial layers in the C1 group (normal IOP and DMSO treatment); (**B-C**) **B** and **C** are the randomly selected magnified views of the yellow and green boxed region in **A** to show representative ZO-1 features.

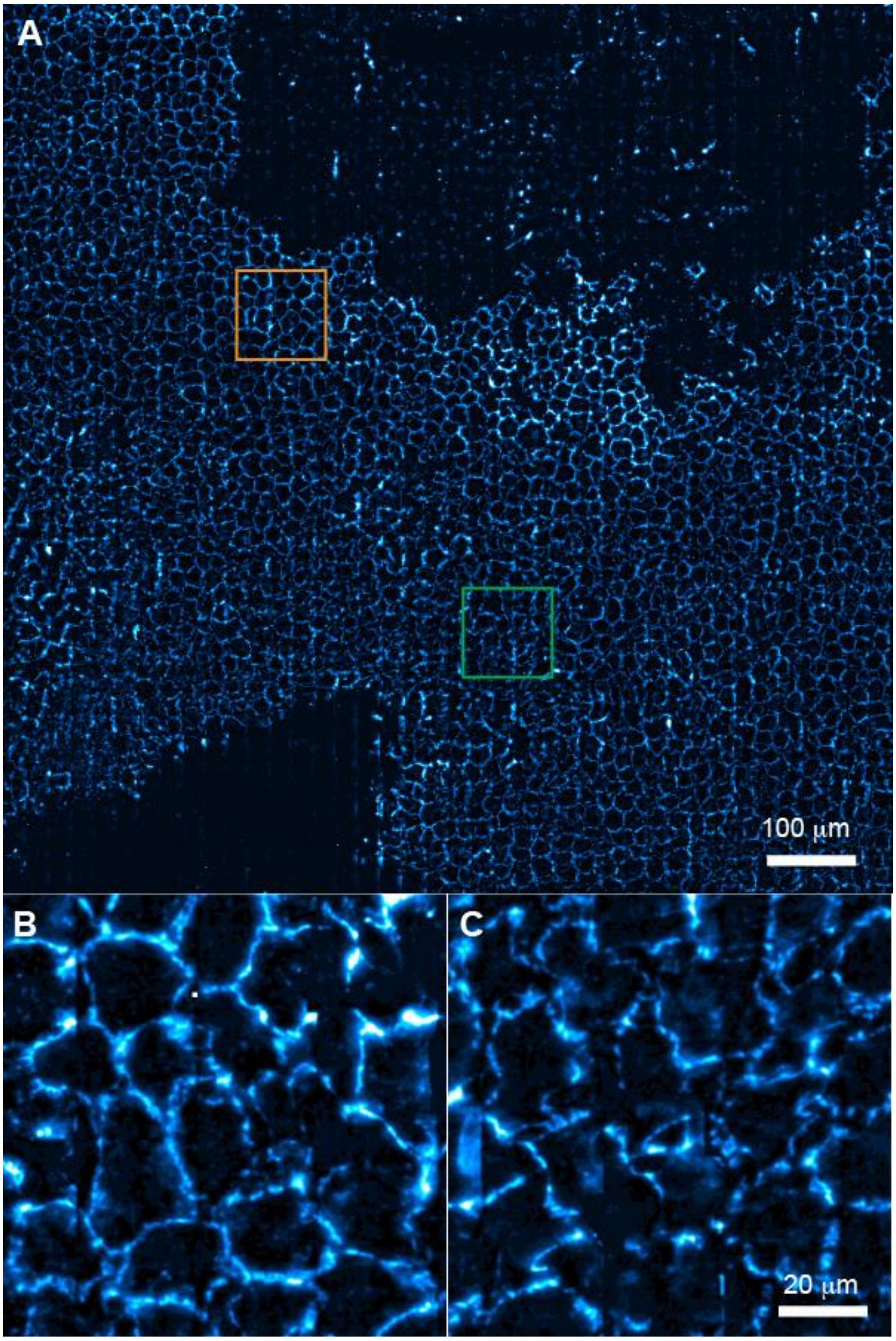


**Fig. S2.** (**A**) A representative tile-scan epifluorescence image with a scanning area of 1x1 $mm^2$ for the corneal endothelial layers in the C2 group (AOH and DMSO treatment); (**B-C**) **B** and **C** are the randomly selected magnified views of the yellow and green boxed region in **A** to show representative ZO-1 features.

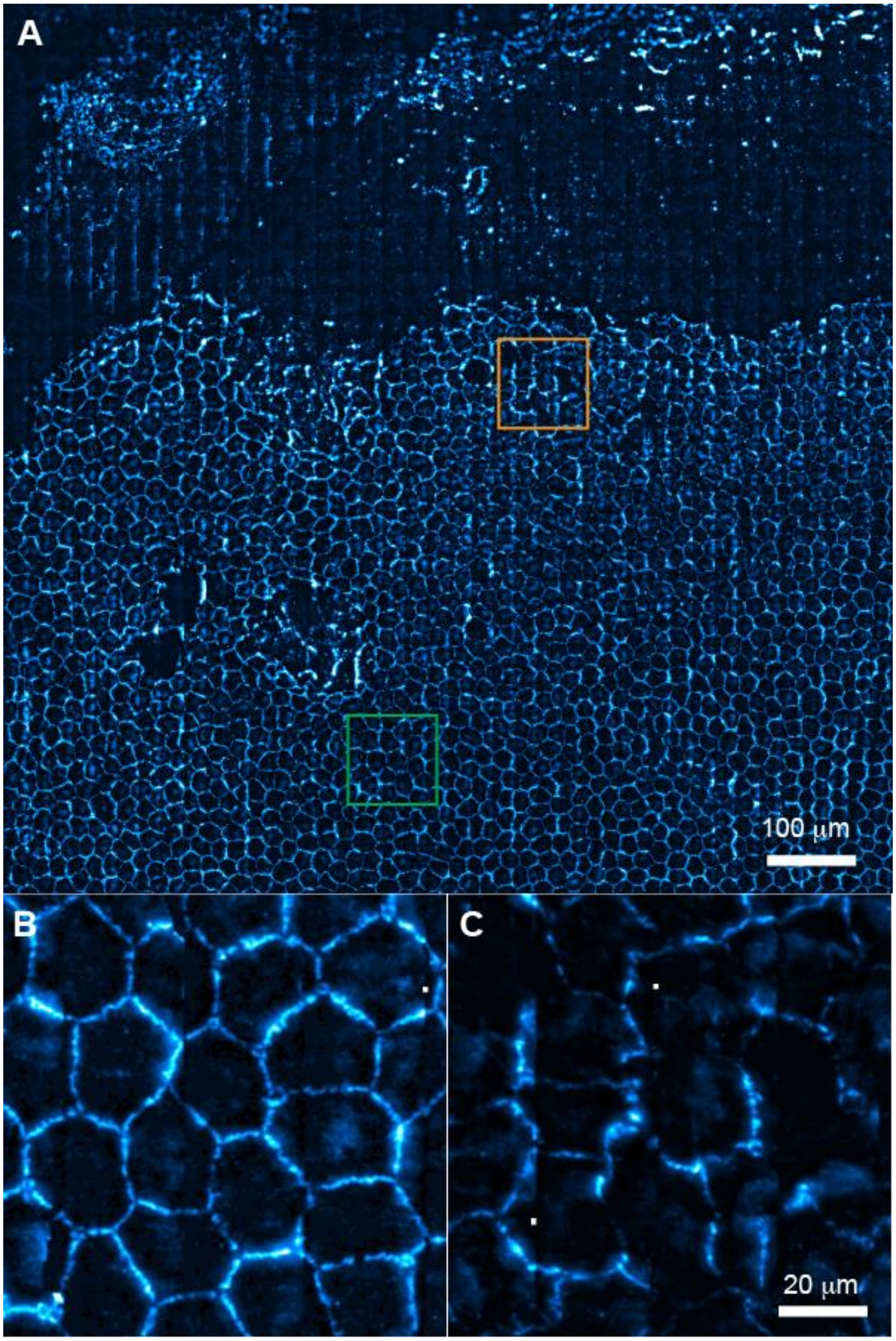


**Fig. S3.** (**A**) A representative tile-scan epifluorescence image with a scanning area of 1x1 $mm^2$ for the corneal endothelial layers in the C3 group (normal IOP and Y-27623 treatment); (**B-C**) **B** and **C** are the randomly selected magnified views of the yellow and green boxed region in **A** to show representative ZO-1 features.

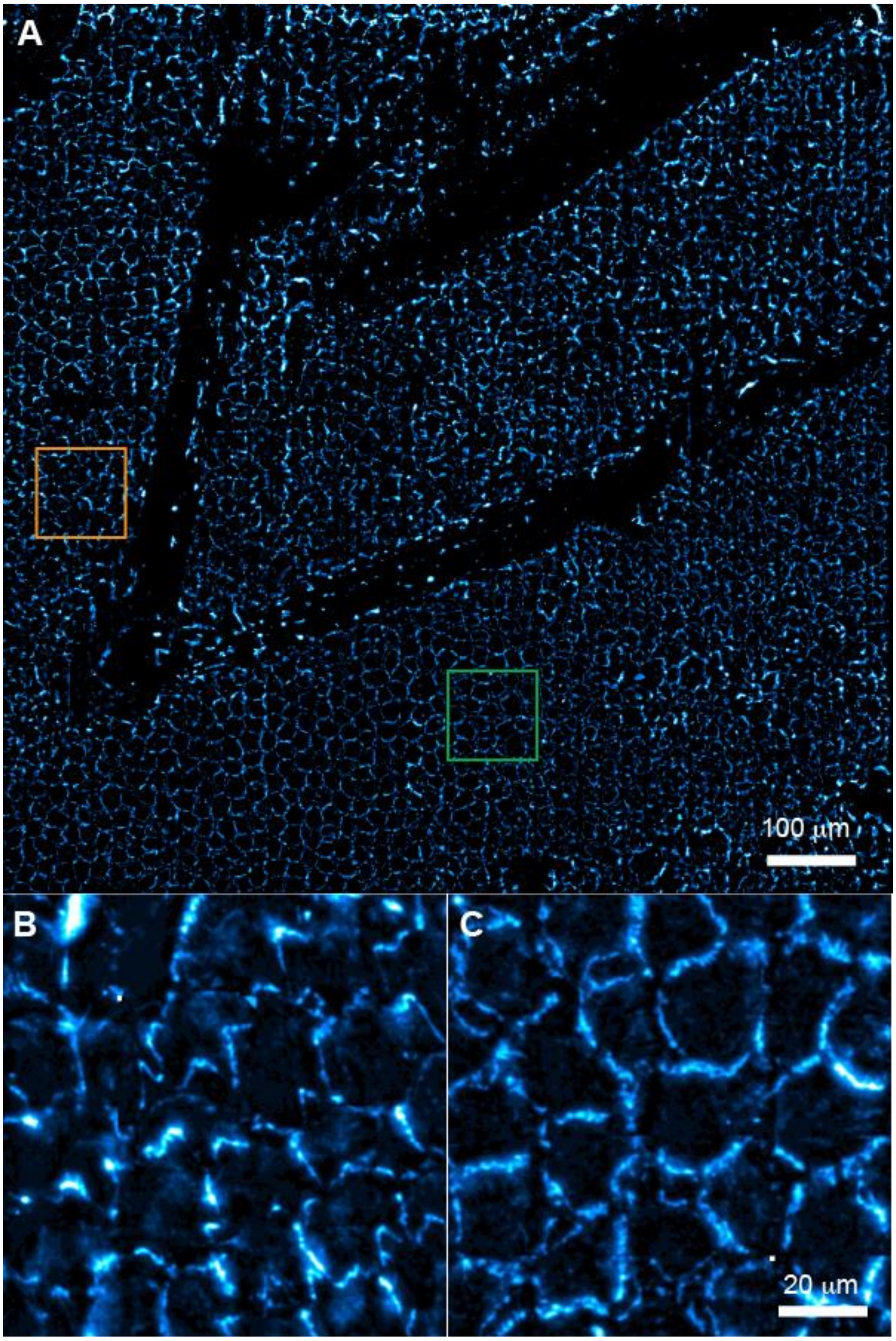


**Fig. S4.** (**A**) A representative tile-scan epifluorescence image with a scanning area of 1x1 $mm^2$ for the corneal endothelial layers in the C4 group (normal IOP and Y-27623 treatment); (**B-C**) **B** and **C** are the randomly selected magnified views of the yellow and green boxed region in **A** to show representative ZO-1 features.
